# DeepIS: deep learning framework for three-dimensional label-free tracking of immunological synapses

**DOI:** 10.1101/539858

**Authors:** Moosung Lee, Young-Ho Lee, Jinyeop Song, Geon Kim, YoungJu Jo, HyunSeok Min, Chan Hyuk Kim, YongKeun Park

## Abstract

The immunological synapse (IS) is a cell-cell junction between T cells and professional antigen presenting cells. Since the IS formation is a critical step for the initiation of an antigen-specific immune response, various live-cell imaging techniques, most of which rely on fluorescence microscopy, have been used to study the dynamics of IS. However, the inherent limitations associated with the fluorescence-based imaging, such as photo-bleaching and photo-toxicity, prevent the long-term assessment of dynamic changes of IS with high frequency. Here, we propose and experimentally validate a label-free, volumetric, and automated assessment method for IS dynamics using a combinational approach of optical diffraction tomography and deep learning-based segmentation. The proposed method enables an automatic and quantitative spatiotemporal analysis of IS kinetics of morphological and biochemical parameters associated with IS dynamics, providing a new option for immunological research.

## Introduction

Understanding the immune response at cellular scale requires knowledge regarding interactions between immune cells and their microenvironment. For T lymphocyte, which is one of the major immune cell types involved in adaptive immune responses, it communicates with antigen-presenting target cells via the formation of a nanoscale cell-cell junction, which is called the immunological synapse (IS). Specifically, the engagement of T cell receptor (TCR) by peptide-loaded MHC complex (pMHC) presented on the target cells leads to the formation of IS that coordinates the downstream signalling events required for the initial activation of T cells (1, 2). Previous studies have shown that the TCR-mediated IS comprises segregated concentric rings of supramolecular activating clusters, which are crucial for stabilization of the IS as well as the secretion of lytic granules (2, 3).

Alternatively, recent studies have highlighted the IS structures formed by chimeric antigen receptor (CAR), a synthetic fusion protein comprised of an extracellular targeting and hinge domain, a transmembrane domain, and intracellular signaling domains (4, 5). The extracellular targeting domain of CAR is typically adopted single-chain variable fragment (scFv) of monoclonal antibodies, allowing the CAR-T cells to recognize various types of surface antigens independent of its MHC restriction. In particular, CD19-specific CART cells have demonstrated remarkable anti-cancer efficacy in patients with B cell malignancies (6). Although CAR/antigen and TCR/pMHC complexes have different IS structures (7), the distinct dynamics and mechanochemical properties of the IS driven by these complexes remain understudied.

A variety of imaging techniques have been used to reveal the hierarchical details of the IS structures and their relevant functions. For instance, electron microscopy and single-molecule localization microscopy have resolved the spatial distributions of subcellular IS compartments beyond optical diffraction limits (8, 9). However, assessing dynamical changes in IS formation requires rapid and continuous imaging of immune cells. Fluorescence microscopy is useful in this regard. It has evolved from total internal reflection fluorescence microscopy for interfacial imaging (10) to light-sheet microscopy for high-speed volumetric imaging (11). Such fluorescence-based techniques have the advantage of chemical specificity. However, they are limited due to photo-bleaching and photo-toxicity, which necessitates the use of complementary label-free, rapid three-dimensional (3D) microscopy methods to assess long-term dynamic changes in IS morphologies (12).

The development of label-free IS imaging has been limited to phase-contrast and differential interference contrast microscopy. The aim is to develop quantitative phase imaging (QPI) as a quantitative label-free imaging method to studying the IS (13). Optical diffraction tomography (ODT) is a promising 3D QPI technique for imaging the 3D refractive index (RI) distribution of cells at a sub-micrometer spatial resolution (14). Unlike nonlinear scanning microscopy that requires a long acquisition time due to weak signal intensities (15-18), ODT enables fast 3D imaging via holographic recording. Also, because the reconstructed RI profile correlates with total cellular protein densities, ODT enables quantitative, photobleaching-free analyses of cell dynamics.

ODT has been actively used to study single-cell morphology (19-22). However, it has not yet been used to study cell-cell interactions that include immune responses. One of the primary reasons is the lack of an accurate 3D segmentation framework to distinguish interacting cell-to-cell interfaces, which is also a problem with other microscopy methods (23). Manual marking is the most primitive segmentation strategy. It is effective but is too laborious and difficult for time-resolved volumetric segmentation. To overcome this barrier, automatic segmentation has been developed based on basic algorithms that include intensity thresholding, filtering, morphological operations, region accumulation, and deformable models (24). However, these methods often result in poor segmentation, particularly for adjoining cell segmentations, which occur in immune responses. To accurately and precisely segment immunologically interacting cells in an automated manner, a novel computational framework is needed.

Here, we present DeepIS, a computational framework for the systematic, label-free analysis of 3D IS dynamics of immune cells in ODT. Our framework is based on deep convolutional neural network (DCNN) that distinguishes adjoining immune cells, target cells, and IS surfaces from the obtained RI tomogram. The proposed framework enables the general, high-throughput, and automated segmentation of more than 1,000 immune-target cell pairs. To validate the method, we applied this method to study the dynamics of CAR/antigen-mediated IS formed between CD19-specific CAR-engineered T cells (CART19) and CD19-positive K562 cancer cells (K562-CD19). The combined use of high-speed imaging capability of ODT enabled 3D high-speed CAR IS tracking in which a tomogram was measured every 3 to 8 seconds for a prolonged period of time (300 seconds to 10 minutes depending on the cell type). Exploiting the linear proportion between RI and protein density(25), we also demonstrate quantitative analyses of CAR IS kinetics by means of the morphological and biochemical properties. The results suggest that DeepIS offers a new analytical approach to immunological research.

## Results

### 3D time-lapse RI measurement of the CART19 and K562-CD19 cell conjugates using optical diffraction tomography

In order to perform ODT experiments in our study, we employed an experimental setup which is based on off-axis holography equipped with a high-speed illumination scanner using a digital micro-mirror device (DMD) (Fig. 1). The setup enables the high-speed acquisition of a single tomogram within 500 milliseconds (Fig. 1a) (26, 27). A 1 × 2 single-mode FC/APC fiber coupler was utilized to split a coherent, monochromatic laser (λ = 532 nm) into sample and reference arms. The DMD was then placed onto the sample plane of the sample arm to control the illumination angle of the first-order diffracted beam striking the sample. To scan the illuminations at large tilt angles, a 4-*f* array consisting of a tube lens (Lens 1, *f* = 250 mm) and a condenser objective (UPLASAPO 60XW, Olympus Inc., Japan) magnified the illumination angle. The light scattered by live cells in a live-cell chamber (TomoChamber, Tomocube Inc., Republic of Korea) was then transmitted through the other 4-*f* array formed by an objective lens (UPLASAPO 60XW, Olympus Inc., Japan) and a tube lens (Lens 2, *f*= 175 mm). The sample beam was combined with the reference beam by a beam splitter and filtered by a linear polarizer. The resultant off-axis hologram was then recorded by a CMOS camera (FL3-U3-13Y3M-C, FLIR Systems, Inc., USA) synchronized with the DMD to record 49 holograms of the sample illuminated at different angles. Using a phase-retrieval algorithm, the amplitude and phase images of the 1:1 conjugate between a CART19 and K562-CD19 cell were retrieved from the measured holograms (Fig. 1b). Based on the Fourier diffraction theorem with Rytov approximation (28, 29), the 3D RI tomogram of the sample was reconstructed from the retrieved amplitude and phase images (Fig. 1c). To fill the uncollected side scattering signals due to the limited numerical apertures of objective lenses, a regularization algorithm based on the non-negative constraint was used (30). The maximum theoretical resolutions of the ODT system were respectively 110 nm laterally and 330 nm axially, according to the Lauer criterion (31). Finally, the reconstructed RI values were converted into corresponding total protein densities using an RI increment of α = 0.185 ml/g (25).

**Figure 1.**
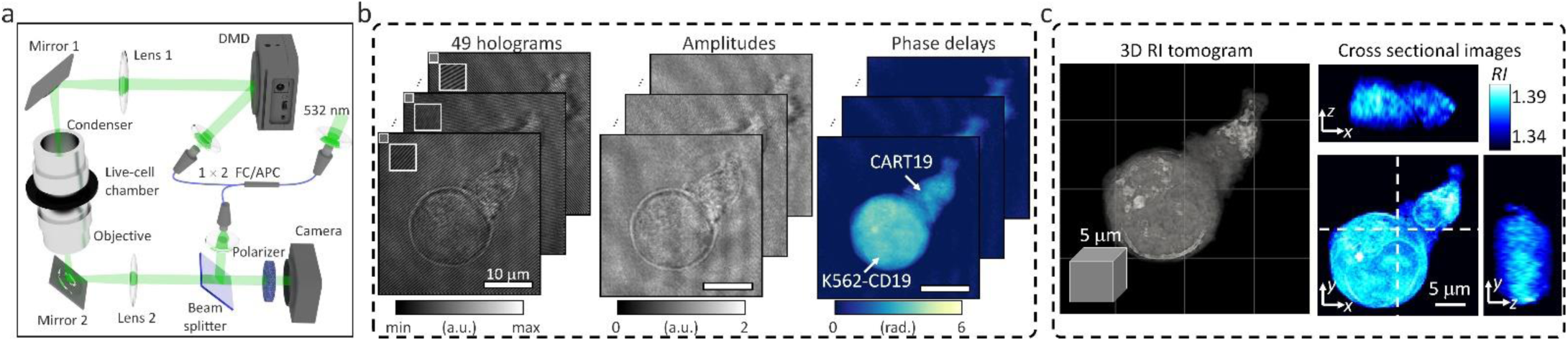
Data acquisition in optical diffraction tomography (ODT). **(a)** The experimental setup for ODT is based on a digital micro-mirror device (DMD) for high-speed illumination scanning. **(b)** Forty-nine holograms of 1:1 conjugate between a CART19 and K562-CD19 cell were recorded at various illumination angles, and their amplitude and phase delay distributions were retrieved. **(c)** A reconstructed refractive index (RI) map.

### DeepIS establishment for automated assessment of CART19 IS dynamics

The IS tracking analysis is preceded by segmentation, which involves dividing volumetric sections for background, cell domains, and IS in the ODT RI map. However, iteration is required for parameter tuning of the manual segmentation method to obtain a single well-segmented label, which is prohibitive to obtain a dynamic dataset. Therefore, we established DeepIS framework based on the DCNN supervised learning method to enable general, high-throughput, and automated segmentation for 3D RI tomograms (Fig. 2). The framework was developed in the following order: (i) Dataset preparation, (ii) Training stage, and (iii) Inference stage. These are detailed next.

**Figure 2.**
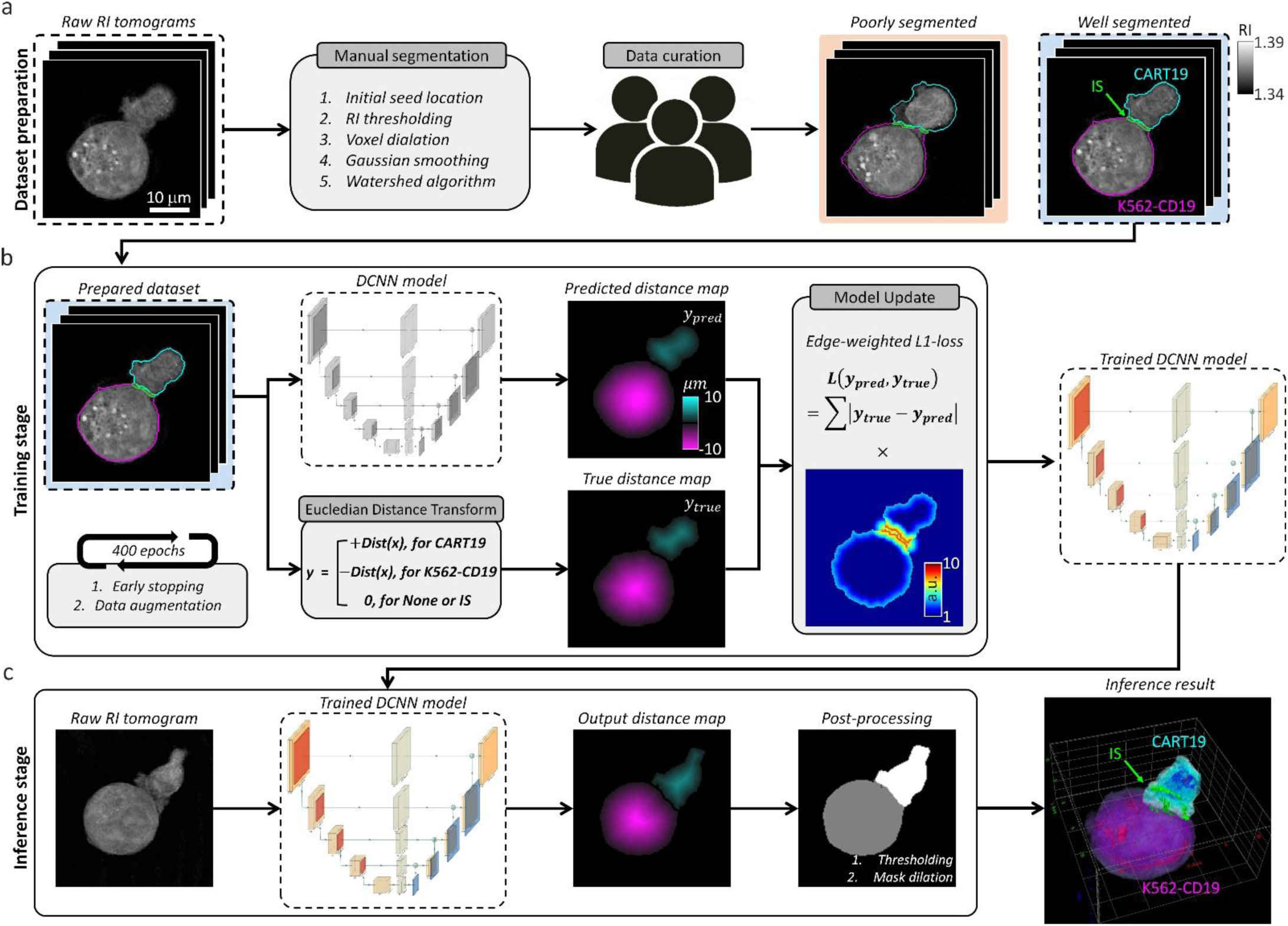
Data flowchart in DeepIS framework. **(a)** Dataset preparation; raw RI tomograms were manually segmented and curated to prepare for training dataset. **(b)** Training stage; the prepared dataset was employed to iteratively train the DCNN model that regresses the distance map of CART19 and K562-CD19 cells with the opposite signs. **(c)** Inference stage; a raw RI tomogram was converted into an output distance map by the trained DCNN model. After post-processing, 3D masks of CART19, K562-CD19, and IS are reconstructed.

#### Dataset preparation

The preliminary step of supervised learning of DCNN is to prepare an annotated dataset (Fig. 2a). To annotate the 3D masks of the CART19 and K562-CD19 cells, we applied a combination of image processing and the watershed algorithm to a raw RI tomogram according to the following steps. First, we manually processed a raw RI tomogram with four hyper-parameters: (i) initial seed locations of each cell to obtain a 3D distance-transform map, (ii) RI threshold for defining cell boundaries, (iii) voxel dilation sizes for merging over-segmented grains into one discrete region, and (iv) standard deviation of the Gaussian smoothing mask. The processed data was then multiplied to the 3D distance-transform map of the cell regions and segmented by the watershed algorithm. Finally, after iterative adjustment of the parameters, experts in cellular biology curated the 236 pairs of well-annotated 3D tomograms obtained from over 2,000 data points. The curated data uniformly reflected the various stages of the IS dynamics, which ensured providing information about the immunological response in both the early and late stages.

#### Training stage

The segmentation tasks were challenged by the lack of distinct boundaries between CART19/K562-CD19 conjugates in RI distributions, diverse morphology of cells, and the demand for precise segmentation at high resolution. As a consequence, although typical segmentation tasks of DCNN were assigned to infer voxel-wise label classification, various types of failure occurred, such as fragmented labels and unnatural IS.

To overcome these limitations and improve the segmentation accuracy and robustness, we designed DCNN to predict the distance map (*y*_*pred*_), which was adapted from a previous study (32) (Fig. 2b, also see Materials and Methods and Supplementary Fig. 1). We first conducted pre-processing of the 3D annotated masks of CART19 and K562-CD19 cells using the Euclidean distance transform to obtain a true distance map (*y*_*true*_). A primary difference from the prior study (32) is that the CART19 and K562-CD19 cells were distinguished by the signs of the distance maps (i.e., positive/negative for CART19/K562-CD19 cells). Presently, we set the background to zero.

With the pre-processed data, the DCNN was optimized using the Adam optimizer (α= 0.5, β = 0.99, initial learning decay value = 0.001) during the training stage to predict the signed 3D distance map of CART19 and K562-CD19 cells. The boundary-weighted L1 function was used as a loss function and indicated the amount of update during the training stage. To prevent overfitting that might arise from a relatively small number of the training data compared with a large number of parameters, early stopping and data augmentation were used. For early stopping, the obtained annotated dataset was into two disjoint subsets. In one subset, 198 tomogram data pairs were used for optimization of model parameters. In the other subset, the remaining 36 tomogram data pairs were used for internal validation. Training of the model parameters was stopped if the performance of the model on the validation set had not improved for five epochs. For data augmentation, a larger annotated dataset was simulated using random rotations, horizontal reflection, cropping, and elastic transformation to make the resulting model more robust to irrelevant sources of variability. The network was trained on four graphics processing units (GPUs; GEFORCE GTX 1080 Ti) for 400 epochs, which took approximately 6 hours. Selection of a model for inference among trained models was based on performance on the internal validation set.

#### Inference stage

In the inference stage, the trained network was used to regress the distance maps of the CART19 and K562-CD19 cells from unlabelled RI tomograms (Fig. 2c). In the post-processing stage, the output distance map was post-processed to yield cell domains masks of each CART19 and K562-CD19 cell through simple thresholding using a value of 54.5 nm, which is approximately half of a voxel pitch. The IS of CART19/K562-CD19 conjugate was defined by dilating the CART19 and K562-CD19 masks and finding an overlapping region.

### Evaluation of DeepIS segmentation performance

Because the label-free segmentation approach aims to distinguish the boundaries between two attached cells at a sub-micrometer spatial resolution, it was essential to validate whether DeepIS provides sufficient segmentation accuracy for our purpose. This was first addressed by comparing the segmentation performances between DeepIS and manual segmentation (Fig. 3). In the trained dataset, this comparison showed that the model could define IS boundaries. Notably, DeepIS generally displayed better segmentation performance than the manual segmentation in the untrained dataset, without notable segmentation problems such as fragmentations and discontinuous boundaries. This observation indicated that DeepIS was well trained and could be exploited to predict the IS boundaries.

**Figure 3.**
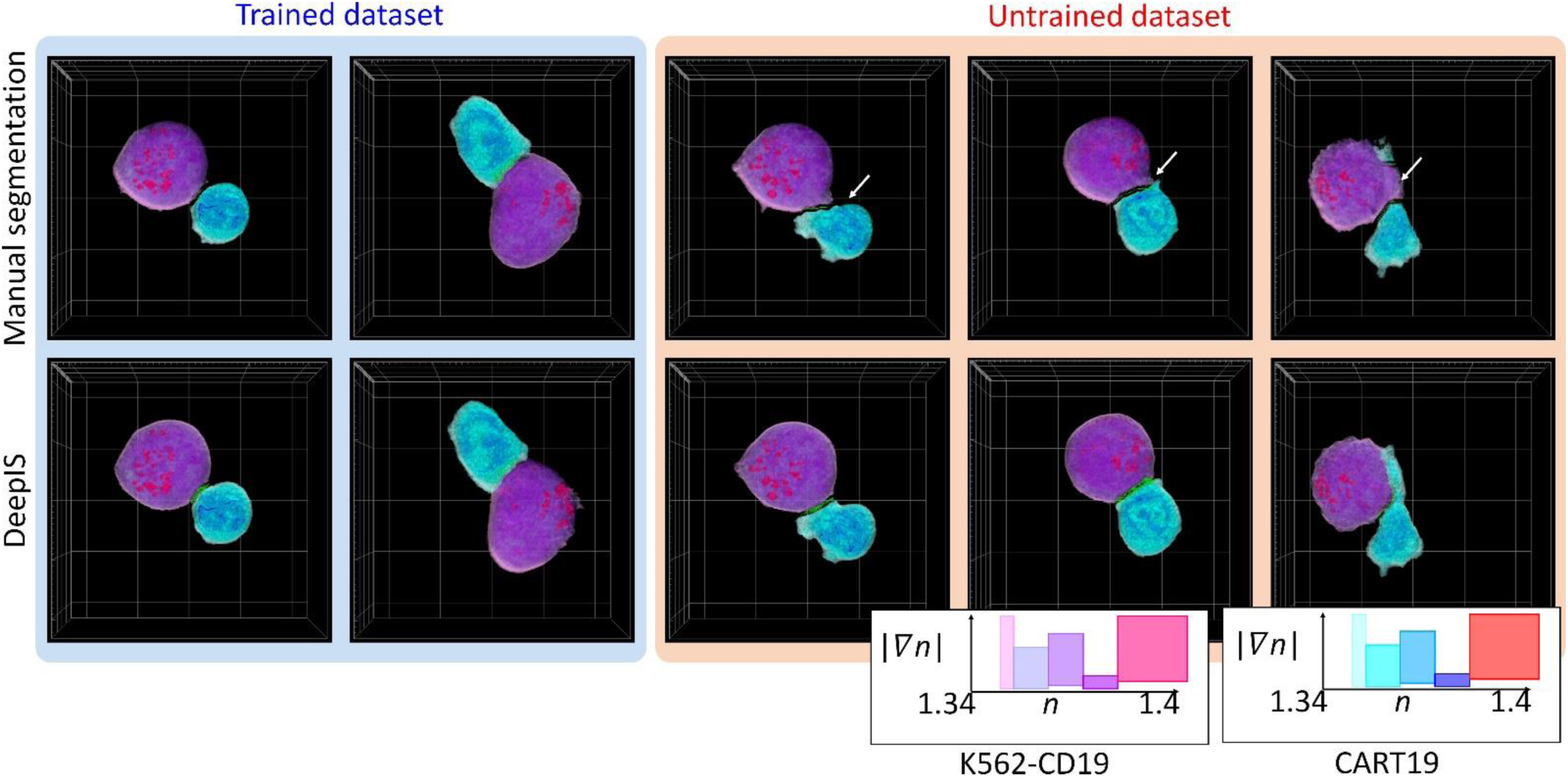
Representative segmentation results. (Top row) Manually segmented masks. (Bottom row) Segmentation results using DeepIS. Blue shade indicates the curated data in the training stage. Red shade indicates the data which were poorly segmented by manual segmentation. White arrows illustrate poorly segmented regions.

**Figure 4.**
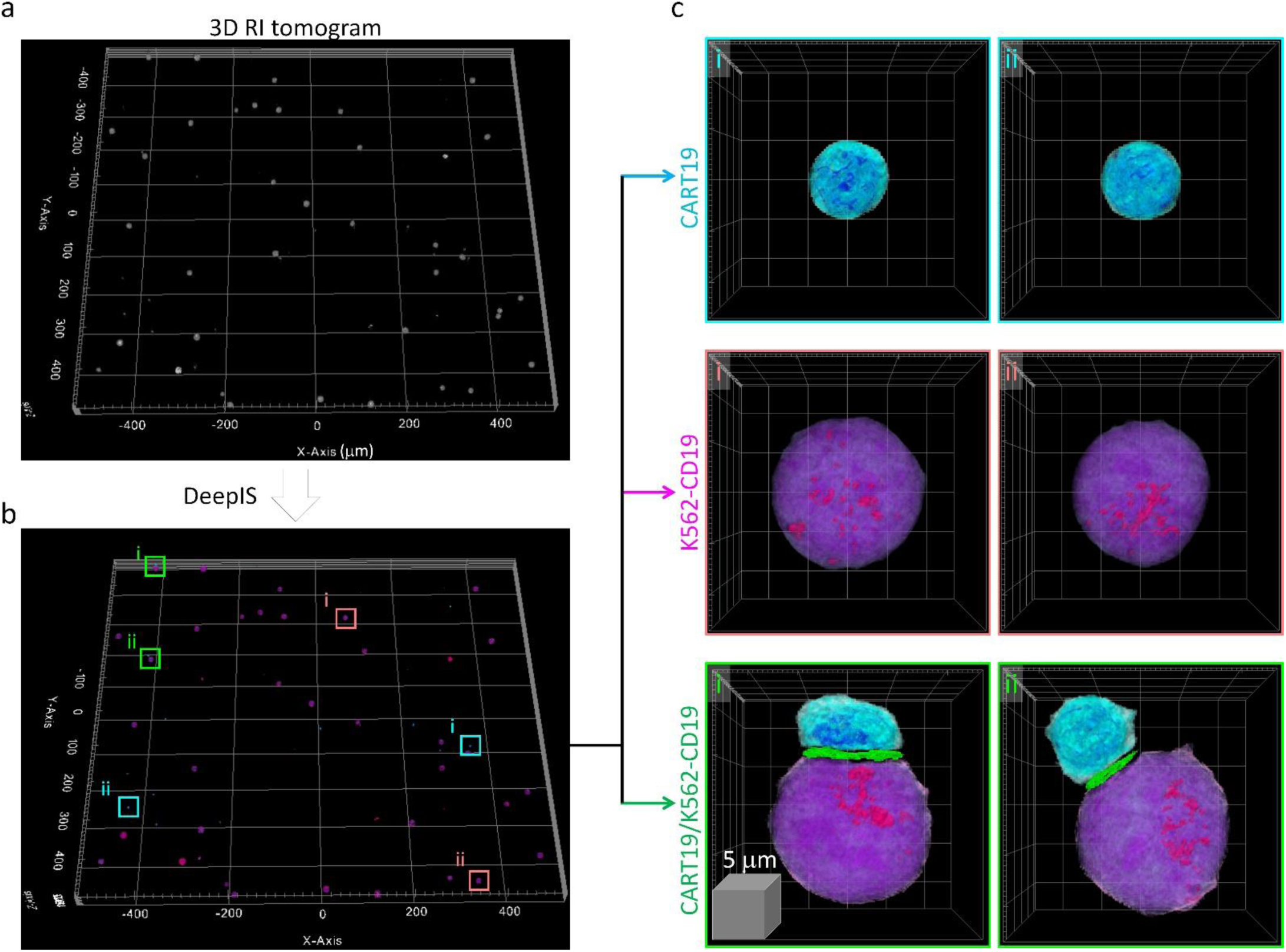
High-throughput semantic segmentation using DeepIS. **(a)** RI tomogram over 0.98 × 1.05 × 0.04 mm^3^ obtained by stitching method. **(b)** Segmentation using DeepIS. **(c)** Representative CART19, K562-CD19, and CART19/K562-CD19 cell conjugates are magnified.

We then examined whether DeepIS could be applied for high-throughput analysis of IS morphology. Wide-area segmentation of CART19 and K562-CD19 cells was carried out over a lateral field-of-view exceeding 1 mm^2^ (Fig. 4a). When cells were located based on RI contrast, DeepIS allowed rapid, automated, and on-site semantic segmentation (Fig. 4b). Interestingly, DeepIS successfully labelled adjoining CART/K562-CD19 cell conjugates as well as individual CART19 and K562-CD19 cells (Fig. 4c), which validated the high-throughput, general segmentation performance of DeepIS.

The segmentation performances of DeepIS were further evaluated by quantifying the segmentation accuracies using manually delineated 3D labels obtained from correlative fluorescence microscopy (Fig. 5). To prepare 3D manual labels, we imaged fixed CART19/K562-CD19 cell conjugates using correlative fluorescence microscopy, and delineated manual labels using the obtained data (Fig. 5a, also see Materials and Methods). The manually drawn labels were compared with the segmentation masks obtained from deepIS using two parameters (Fig. 5b). We first quantified 3D intersection-over-union for volumetric masks. The mean ± standard deviation (SD) values of CART19 and K562-CD19 labels were 0.866 ± 0.040 and 0.860 ± 0.082 respectively, implying a greater than 80% good overlap between the manual labels and automatically segmented labels using DeepIS. The segmentation accuracy of IS labels was evaluated by quantifying boundary displacement error. The mean ± SD value was 3.82 ± 2.09 voxels, which corresponded to a sub-micrometer displacement error of 832.9 ± 457.2 nm. These quantitative analyses supported the conclusion that the deep learning model could be generally utilized throughout the IS analyses with high fidelity.

**Figure 5.**
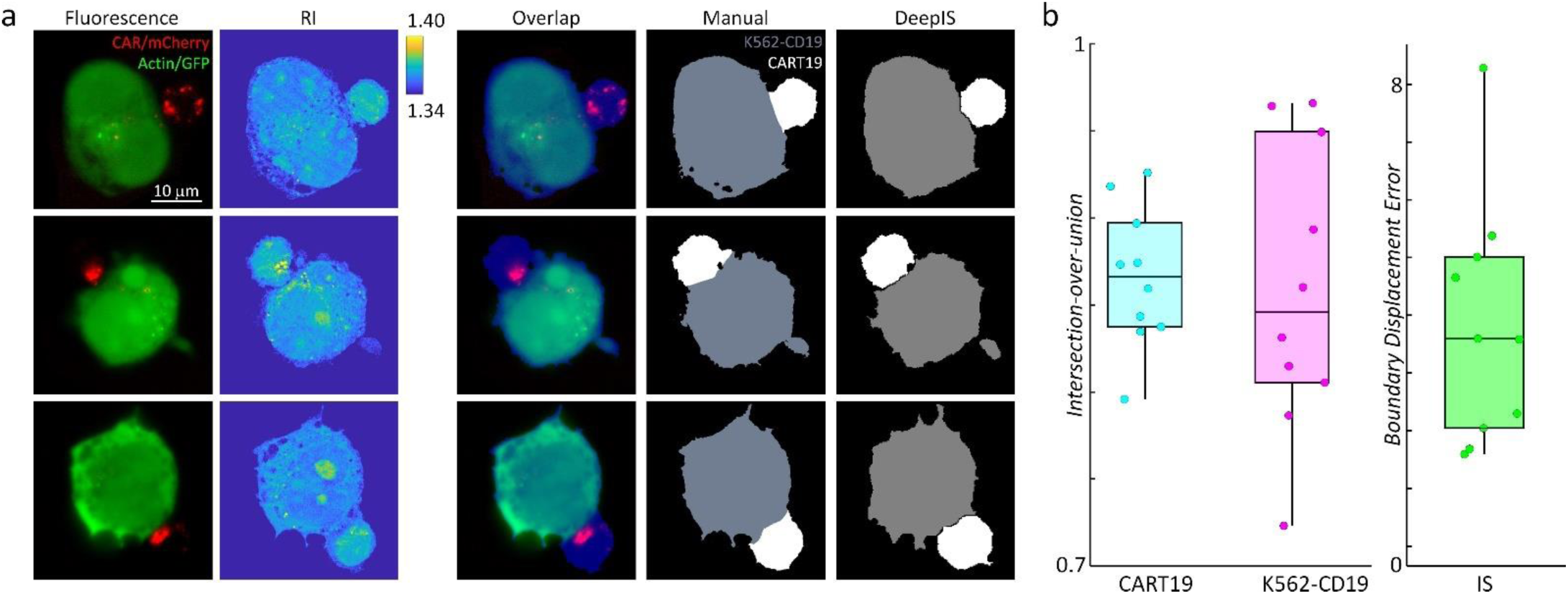
Quantitative analyses of segmentation performances using correlative fluorescence microscopy and ODT. **(a)** Representative fluorescence sections (first column), RI sections (second column), and their overlapped images (third column). CART19 and K562-CD19 labels were manually delineated based on the correlative images (fourth column) and compared with the labels obtained from DeepIS (fifth column). **(b)** Quantifications of DeepIS segmentation performances (*n* = 10). Intersection-over-union was measured for CART19 (0.866 ± 0.040) and K562-CD19 (0.860 ± 0.082). Boundary displacement error was measured for IS (3.816 ± 2.09 voxels; 832.9 ± 457.2 nm). Each boxplot indicates the median, upper, and lower quartiles of each population.

### Quantitative kinetic analysis of CART19 IS formation using DeepIS

The successfully established DeepIS was implemented in the detailed kinetic analysis of the IS formation between CART19 and K562-CD19 cells by means of morphological and biochemical parameters (Fig. 6). Specifically, we analyzed 27 sets of IS dynamic datasets measured over 300 seconds to 10 minutes at time intervals of 3 to 8 seconds to determine the kinetics of synapse area, membrane protein density, and intracellular protein density.

**Figure 6.**
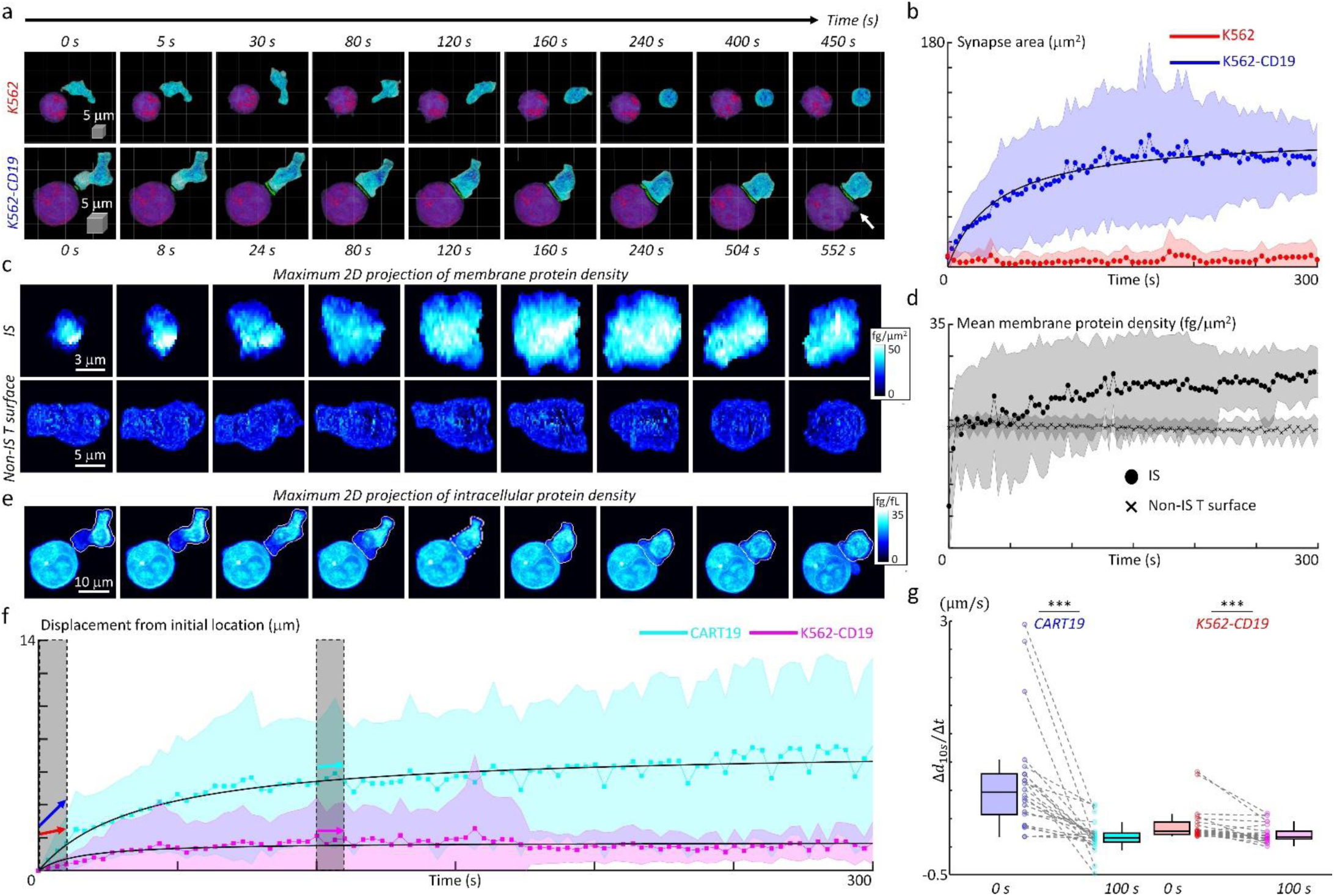
Quantification of initial IS formation kinetics of CART19 cells. **(a)** Representative snapshots of a video of CART19 cells responding to K562 (top row) and K562-CD19 (bottom row) cells. Purple : K562, Blue : CART19. The 0-second indicates the initial contact point of the effector cell and target cells. The white arrow indicates the blebbing point. **(b)** Temporal changes in the synapse areas of CART19 cells responding to K562 cells (*n* = 5) and K562-CD19 cells (n = 22). Black line: fitting curve with *A*(*t*) = *A*_*max*_*t*/(*t* + *τ*_1/2_), where *A*_*max*_ = 106.16 μm^2^ and *τ*_1/2_ = 39.63 seconds, Pearson correlation coefficient = 0.93. **(c**) Maximum 2D projection of membrane protein density of IS (top row) and non-IS T surface (bottom row) for CART19/K562-CD19 conjugate. **(d)** Temporal changes in the mean membrane protein densities of CART19 immunological synapses (circles) and non-IS T surfaces (crosses) responding to K562-CD19 cells. The mean membrane protein densities of IS and non-IS T surface at 300 seconds were 27 ± 4 and 19 ± 1 fg/μm^2^, respectively. **(e)** Maximum 2D projection of intracellular protein density distributions of the CART19/K562-CD19 conjugate. The white contours indicate the boundaries of the CART19 mask. **(f)** Temporal changes in the displacements of the center-of-masses of CART19 and K562-CD19 cells from their initial locations. Black lines : Δ*d*(*t*) = Δ*d*_max_ *t*/(*t* + *τ*_1/2_), where Δ*d*_*max*_ = 7.48 and 1.74 μm, *τ*_1/2_ = 37.84 and 14.56 seconds, Pearson correlation coefficient = 0.91 and 0.58 for CART19 and K562-CD19 respectively. **(g)** The average changes for 10 seconds in the early (0 seconds) and late (100 seconds) stages are marked by colored arrows, and statistically compared by two-tail paired Wilcoxon tests for CART19 and K562-CD19 cells, respectively. CART19: Δ*d/*Δ*t*_*10s*_= 818.5 ± 788.5 nm/s at 0 seconds, and 26.6 ± 223.6 nm/s at 100 seconds. K562-CD19: Δ*d/*Δ*t*_*10s*_= 194.3 ± 246.5 nm/s at 0 seconds, and 63.7 ± 125.6 nm/s at 100 seconds. *** indicates *p* < 0.001 for CART19 and K562-CD19. Each boxplot indicates the median, upper, and lower quartiles of each population. The central points and shades in Figs. 6b, 6d, 6f, 6g indicate the mean and standard deviations respectively.

First, we examined the temporal changes of synapse areas depending on the expression of the target antigen, CD19, on K562 cells (Fig. 6a). As expected, CART19 cells could not form a stable synapse with K562 cells (CD19-negative) in five independent experimental trials (Fig. 6a, Supplementary Video 1). By contrast, CART19 cells formed a stable IS with K562-CD19 cells (CD19-positive), and induced apoptotic blebbing on the target cells about 9 minutes after the initial contact (Supplementary Video 2). For the statistical analysis of the initial IS area changes, DeepIS was applied to both dynamic datasets and successfully segmented CART cells, target cells, and IS boundaries. The temporal graphs of the mean synapse area showed that, whereas the IS for K562 cells was not stably formed for 300 seconds, the IS for K562-CD19 expanded to the half of the maximum synapse area (*A*_*max*_ = 106.16 μm^2^) within 40 seconds and reached a steady state within only 3 minutes (Fig. 6b).

Next, we also assessed whether the membrane protein amount differs between the IS and non-IS areas of CART19 cells by comparing the 2D maximum projected snapshots of membrane protein densities in each area (Fig. 6c). Within the IS surface, a dramatic increase in membrane protein density as well as the synapse area was observed. By contrast, the non-IS T surfaces of CART19 cells maintained the constant, lower membrane protein densities. In line with this observation, quantification of temporal changes in the mean membrane protein densities revealed that up to 27 ± 4 fg/μm^2^ at 300 seconds have been accumulated in the IS surface, which was higher than average protein density (19 ± 1 fg/μm^2^) in the non-IS surface (Fig. 6d).

We further explored the cell mechanics of CART19 and K562-CD19 cells during their initial IS formation. As in Fig. 6e, the time-lapse snapshots of maximum 2D projection of intracellular protein densities visualized the dynamic action of CART19 cells, which incorporated the polarization of the intracellular protein contents. Furthermore, CART19 cells exerted mechanical forces during the dynamic IS formation, which led to the subsequent translational displacement and deformation of the K562-CD19 cells (Supplementary Video 3). To quantify the cell translocation dynamics, we traced the temporal changes in the displacement of the center-of-mass (the average displacement weighted by the intracellular protein density) with respect to the initial location for each cell (Fig. 6f). As observed, CART19 cells exhibited more dynamic motions (Δ*d*_*max*_ = 7.48 μm) than K562-CD19 cells (Δ*d*_*max*_ = 1.74 μm) during the IS formation. For both cells, the mechanical translocations were more dramatic in the earlier stage of the IS formation, as confirmed statistically by comparing the average change in the cell translocations in the early and late stage (Fig. 6g).

Overall, the results presented here generally recapitulate previously observed phenomena in the synapse studies based on conventional microscopy techniques. The rapid increase in the membrane protein density in the synapse area likely reflects the influx of large amounts of IS protein components, including CARs, actins, and other adhesion molecules (33). In addition, perhaps the intracellular protein density changes in CART19 cells, which indicates polarization of the intracellular organelles in CART19 cells until stabilization, can be explained by the centrosome polarization of cytotoxic T lymphocytes (34). Lastly, the force exerted by the CART19 cells on target cells during the IS formation suggests that CART cells also integrate mechanical potentiation during target cell killing similar to TCR-based cytotoxic T cells (35).

## Discussion

The collective results demonstrate that DeepIS in combination with ODT and deep neural network enables the label-free, time-lapse 3D IS tracking in an automated manner. The successful segmentation performances of DeepIS could not be possible without careful validations in the initial development stage, including data curation, model training, and qualitative and quantitative tests for general segmentation capabilities. This platform was applied to define the IS parameters related to their morphological and biochemical traits, and to quantify the IS dynamics of CART19 cells. cells. We anticipate that the proposed method can be generalized to study a broad range of IS that are mediated by different types of immune receptors such as TCR and B cell receptor (BCR) as well as invariant receptors expressed on innate natural killer (NK) cells. In particular, the model will be a powerful way to test whether the IS morphology is affected by chemical treatment and genetic mutations (36).

The present work is, to the best of our knowledge, the first application of ODT and deep learning to understand IS dynamics. Refinements are anticipated in an immediate follow-up study. For example, more rapid immune dynamics can be investigated at higher spatiotemporal resolution in a more sophisticated experimental setup, which may reveal dynamic diffusions of subcellular organelles (37) during the immune responses. Also, more efforts in data curation and network design may allow robust time-lapse IS tracking. In order to do so, segmentation based on active contour methods, and new network models based on recurrent neural networks (38), Bayesian neural network (39), and pyramid pooling (40) may be incorporated.

Our study focused on the platform capable of quantitatively tracking 3D IS dynamics in a completely label-free manner. The RI contrast used in the present study alone does not provide information about individual proteins that are transported during the IS formation of CART19 cells. To provide such biochemically relevant information, we expect that correlative imaging with fluorescence microscopy will circumvent the inherent lack of chemical specificities (41-43). It also remains unclear how much force was associated during the IS formation, and whether this force correlates with the cytotoxic intensity of CART19 cells. To test the hypothesis, the simultaneous use of microscopic force measurement techniques and ODT will be helpful. For instance, a combined technique of ODT and holographic optical tweezers (44, 45) or traction force microscopy can supplement our DeepIS framework for understanding the mechanical potentiation during the IS kinetics. We expect that these future studies may be exploited for label-free predictions of dynamic molecular transports occurring in the IS formation, and provide complementary information required to elucidate the IS formation mechanisms.

### Significance statement

Rapid, label-free, volumetric, and automated assessment in microscopy is necessary to understand the dynamic interactions between immune T cells and their targets through the immunological synapse (IS) and the relevant immunological functions. However, attems to realize the automatic tracking of IS dynamics have been stymided by limitations of imaging techniques and computational analaysis methods. Here, we demonstrate the 3D IS tracking by combining optical diffraction tomography and deep-learning-based segmentation. The proposed approach enables automatic and quantitative spatiotemporal analyses of IS kinetics regarding morphological and biochemical parameters related to the protein densities of immune cells. As a complementary method to fluorescence microscopy, the proposed method will offer a new perspective for studies in immunology.

## Materials and Methods

*SI Appendix* contains architecture design of our deep neural network. The raw model output data and codes are available from the authors upon request.

### Cell preparation and establishment of cell lines

#### Cell lines and culture

K562 cells and CD19-positive K562 cells (K562-CD19; target cells) were kindly provided by Travis S. Young (California Institute for Biomedical Research). The cells were cultured in RPMI-1640 medium supplemented with 10% heat-inactivated fetal bovine serum (FBS), 2 mM L-glutamine, and 1% penicillin/streptomycin in a humidified incubator with a 5% CO_2_ atmosphere at 37°C. The Lenti-X™ 293T cell line was purchased from Takara Bio (Japan). The cells were maintained in Dulbecco’s modified Eagle medium supplemented with 10% heat-inactivated FBS, 2 mM L-glutamine, 0.1 mM non-essential amino acids, 1 mM sodium pyruvate, and 1% penicillin/streptomycin.

#### Plasmid construction

CD19-specific chimeric antigen receptor (CD19-BBz CAR) was synthesized. The construct is composed of anti-CD19 scFv (FMC63) connected to a CD8α spacer domain and CD8α transmembrane domain, 4-1BB (CD137) co-stimulatory domains, and the CD3ζ signaling domain (46). The cytoplasmic domain comprised of a truncated CD271(ΔLNGFR) gene for the isolation of T cells expressing CAR was amplified from pMACS-ΔLNGFR (Miltenyi Biotec, Germany) and overlapped with the P2A oligonucleotide. The ΔLNGFR-P2A gene was overlapped with the CD19-BBz CAR gene and then inserted into the BamHI and SalI sites of pLV vectors to generate pLV-ΔLNGFR-P2A-CD19-BBz CAR.

To define the IS of CD19-specific CAR-T (CART19; effector) cells, we generated a mCherry-tagged CD19-BBz CAR. The mCherry gene was amplified from pLV-EF1a-MCS-IRES-RFP-Puro (Biosettia, USA) and overlapped with synthetic oligonucleotides of a G4S linker. The G4S-mCherry gene was inserted at the 3`-end of CD19-BBz CAR to generate pLV-BBz-CAR-mCherry.

#### Generation of CAR-transduced human T cells

To generate a recombinant lentivirus supernatant, 6×10^5^ Lenti-X™ 293T cells were cultured in wells of a six-well plate for 24 hours and then transfected with the lentivirus packaging vectors (pMDL, pRev, pMDG.1) and the pLV vectors encoding ΔLNGFR-P2A-CD19-BBz CAR or BBz-CAR-mCherry using 10 μL of Lipofectamine2000 (Thermo Fisher Scientific, USA). Two days after transfection, the lentivirus containing supernatant was collected and stored at −80°C until used.

Peripheral blood mononuclear cells (PBMCs) were separated from whole blood samples of healthy donors using SepMate™ tubes (STEMCELL Technologies, Canada) following the manufacturer’s instructions. The PBMCs were stimulated with 4 μg/mL of plate-bound anti-CD3 antibody (clone OKT3; Bio X cell), 2 μg/mL of soluble anti-CD28 antibody (clone CD28.2; Bio X cell), and 300 IU/mL human recombinant IL-2 (BMI KOREA, Republic of Korea).

Two days after stimulation, the activated T cells were mixed with the lentivirus supernatant, centrifuged at 1000×*g* for 90 minutes, and incubated overnight at 37°C. CAR-transduced T cells were cultured at a density of 1×10^6^ cells/mL in RPMI-1640 supplemented with 10% heat-inactivated FBS, 2 mM L-glutamine, 0.1 mM non-essential amino acid, 1 mM sodium pyruvate, and 55 μM β–mercaptoethanol in the presence of human recombinant interleukin (IL)-2 (300 IU/mL) until isolation of CAR-expressing T cells from bulk T cells.

#### Flow cytometry

The percentage of CAR and ΔLNGFR positive T cells was assessed by biotin-conjugated rhCD19-Fc (Cat # CD9-H5259, ACRO Biosystems, USA) with AF647-conjugated streptavidin (Cat # 405237, Biolegend, USA), and fluorescein isothiocyanate (FITC)-conjugated LNGFR antibody (Cat# 130-112-605, Miltenyi Biotec, Germany).

#### Isolation of CAR-transduced T cells

CAR-and ΔLNGFR-positive T cells were isolated using the human CD271 MicroBead kit (Cat# 130-099-023, Miltenyi Biotec) following the manufacturer’s instructions. Sorted CART19 cells were expanded for 6 days with RPMI-1640 medium supplemented with 10% heat-inactivated FBS, 2 mM L-glutamine, 0.1 mM non-essential amino acids, 1 mM sodium pyruvate, and 55 μM β–mercaptoethanol in the presence of recombinant human (rh)IL-2 (300 IU/mL). For isolation of CD4^+^ or CD8^+^ CART19 cells, CART19 cells were stained with FITC-conjugated anti-CD4 (Cat# 11-0048-41, Thermo Fisher Scientific) and PerCP-Cy5.5-conjugated anti-CD8 antibody (Cat# 344710, Biolegend). CD4^+^ or CD8^+^ CART19 cells were sorted up to 98% purity using the MoFlo Astrios sorter (Beckman Coulter, USA). Likewise, mCherry expressing CART19 cells also isolated using the MoFlo Astrios sorter (Beckman Coulter).

### Correlative fluorescence microscopy

To evaluate the segmentation performances quantitatively, the evaluation data were prepared using correlative imaging between wide-field fluorescence microscopy and ODT (HT-2H, Tomocube Inc., Republic of Korea) (43). CART19 and K562-CD19 cells were imaged in different fluorescence channels by tagging CAR of the CART19 cells and actin of the K562-CD19 cells with mCherry and GFP protein, respectively. The fluorescence-tagged cells were fixed by 4% paraformaldehyde solution, and the remaining solution was replaced with fresh DMEM solution. To excite mCherry and GFP proteins of the prepared cells, blue and green light-emitting diodes were illuminated in wide-field epi-fluorescence geometry. Sixty-three 3D fluorescence image stacks were obtained by scanning the objective lens with an axial spacing of 313 nm. The obtained two-channel images were deconvolved using the blind Lucy algorithm with theoretical 3D point spread functions as initial estimates. The ground-truth labels of the CART19 and K562-CD19 cells were derived from expert biologists who manually thresholded, delineated, and smoothed the cell volume by means of the overlapped RI and fluorescence images.

### Design of DCNN architecture

The DCNN architecture was designed on the basis of UNet architecture, which features excellent performance for various biomedical volumetric segmentation tasks such as multi-cell(47), organ(48), and tumor segmentation tasks(49). Our model employed five contracting and expanding layers comprising 32, 64, 128, 256, and 512 filters, respectively. To improve the segmentation performance, several modifications of the architecture were added while maintaining the overall U-shaped feature map flow line. First, we employed a series of ResNet blocks from ResNet(50) in the contracting paths for extracting the feature more robustly. Also, to increase the receptive field, we employed the feature skip connection that passes through the global convolutional network layer(51) with *k* = 13, 13, 9, 7, and 5, respectively. The overall schematic figure of DCNN architecture is shown in Supplementary. Fig. 1. Our network was implemented in Python using the PyTorch package (http://pytorch.org), and the processing steps were performed in MATLAB (MathWorks, Inc.).

### Statistical Analysis

MATLAB was used in order to calculate P values by two-tail paired Wilcoxon tests to compare the sample means in Fig. 6h. All of the numbers following the ± sign in the text are standard deviations.

## Acknowledgments

This work was supported by KAIST, Tomocube Inc., National Research Foundation of Korea (NRF) (2017M3C1A3013923, 2015R1A3A2066550, 2018K000396, 2018R1A6A3A01011043), the Bio & Medical Technology Development Program of NRF funded by the Ministry of Science & ICT (2014M3A9D8032525), KAIST GCORE(Global Center for Open Research with Enterprise) grant funded by the Ministry of Science and ICT (N11190028).

## AUTHOR CONTRIBUTIONS

Y.-K.P. and C.-H.K. initiated the work and supervised the project. M.L. and Y.-H.L. performed the experiments. M.L. and J.S. developed the methods. All authors wrote the manuscript.

## References

1. Bromley SK, et al. (2001) The Immunological Synapse. Annu. Rev. Immunol. 19(1):375–396.

2. Basu R & Huse M (2017) Mechanical Communication at the Immunological Synapse. Trends in Cell Biology 27(4):241–254.

3. Monks CRF, Freiberg BA, Kupfer H, Sciaky N, & Kupfer A (1998) Three-dimensional segregation of supramolecular activation clusters in T cells. Nature 395:82.

4. Lee Y-H & Kim CH (2019) Evolution of chimeric antigen receptor (CAR) T cell therapy: current status and future perspectives. Archives of Pharmacal Research:1–10.

5. Mukherjee M, Mace EM, Carisey AF, Ahmed N, & Orange JS (2017) Quantitative Imaging Approaches to Study the CAR Immunological Synapse. Mol Ther 25(8):1757–1768.

6. van der Stegen SJ, Hamieh M, & Sadelain M (2015) The pharmacology of second-generation chimeric antigen receptors. Nat Rev Drug Discov 14(7):499–509.

7. Davenport AJ, et al. (2018) Chimeric antigen receptor T cells form nonclassical and potent immune synapses driving rapid cytotoxicity. Proc Natl Acad Sci U S A 115(9):E2068–E2076.

8. Choudhuri K, et al. (2014) Polarized release of T-cell-receptor-enriched microvesicles at the immunological synapse. Nature 507:118.

9. Hu YS, Cang H, & Lillemeier BF (2016) Superresolution imaging reveals nanometer-and micrometer-scale spatial distributions of T-cell receptors in lymph nodes. Proceedings of the National Academy of Sciences 113(26):7201.

10. Balagopalan L, Sherman E, Barr VA, & Samelson LE (2010) Imaging techniques for assaying lymphocyte activation in action. Nature Reviews Immunology 11:21.

11. Chen B-C, et al. (2014) Lattice light-sheet microscopy: Imaging molecules to embryos at high spatiotemporal resolution. Science 346(6208):1257998.

12. Skylaki S, Hilsenbeck O, & Schroeder T (2016) Challenges in long-term imaging and quantification of single-cell dynamics. Nat Biotechnol 34(11):1137–1144.

13. Park Y, Depeursinge C, & Popescu G (2018) Quantitative phase imaging in biomedicine. Nature Photonics 12(10):578–589.

14. Kim K, et al. (2016) Optical diffraction tomography techniques for the study of cell pathophysiology. Journal of Biomedical Photonics and Engineering 2(2).

15. Zumbusch A, Holtom GR, & Xie XS (1999) Three-Dimensional Vibrational Imaging by Coherent Anti-Stokes Raman Scattering. Physical Review Letters 82(20):4142–4145.

16. Freudiger CW, et al. (2008) Label-Free Biomedical Imaging with High Sensitivity by Stimulated Raman Scattering Microscopy. Science 322(5909):1857.

17. Squier JA, Müller M, Brakenhoff GJ, & Wilson KR (1998) Third harmonic generation microscopy. Opt. Express 3(9):315–324.

18. Zipfel WR, et al. (2003) Live tissue intrinsic emission microscopy using multiphoton-excited native fluorescence and second harmonic generation. Proceedings of the National Academy of Sciences 100(12):7075.

19. Yoon J, et al. (2017) Identification of non-activated lymphocytes using three-dimensional refractive index tomography and machine learning. Scientific Reports 7(1):6654.

20. Kim G, Lee S, Shin S, & Park Y (2018) Three-dimensional label-free imaging and analysis of Pinus pollen grains using optical diffraction tomography. Scientific Reports 8(1):1782.

21. Park Y, et al. (2008) Refractive index maps and membrane dynamics of human red blood cells parasitized by *Plasmodium falciparum*. Proceedings of the National Academy of Sciences 105(37):13730–13735.

22. Yang S-A, Yoon J, Kim K, & Park Y (2017) Measurements of morphological and biophysical alterations in individual neuron cells associated with early neurotoxic effects in Parkinson’s disease. Cytometry Part A 91(5):510–518.

23. Uchida S (2013) Image processing and recognition for biological images. Development, Growth & Differentiation 55(4):523–549.

24. Mayer CE, Rudolf F, Stelling J, & Dimopoulos S (2014) Accurate cell segmentation in microscopy images using membrane patterns. Bioinformatics 30(18):2644–2651.

25. Barer R, Ross KF, & Tkaczyk S (1953) Refractometry of living cells. Nature 171(4356):720–724.

26. Shin S, Kim K, Yoon J, & Park Y (2015) Active illumination using a digital micromirror device for quantitative phase imaging. Opt Lett 40(22):5407–5410.

27. Lee K, Kim K, Kim G, Shin S, & Park Y (2017) Time-multiplexed structured illumination using a DMD for optical diffraction tomography. Opt Lett 42(5):999–1002.

28. Wolf E (1969) Three-dimensional structure determination of semi-transparent objects from holographic data. Optics Communications 1(4):153–156.

29. Devaney AJ (1981) Inverse-scattering theory within the Rytov approximation. Opt Lett 6(8):374–376.

30. Lim J, et al. (2015) Comparative study of iterative reconstruction algorithms for missing cone problems in optical diffraction tomography. Opt. Express 23(13):16933–16948.

31. Lauer V (2002) New approach to optical diffraction tomography yielding a vector equation of diffraction tomography and a novel tomographic microscope. J Microsc 205(2):165–176.

32. Wang W, et al. (2018) Learn to segment single cells with deep distance estimator and deep cell detector. eprint arXiv:1803.10829:arXiv:1803.10829.

33. Xiong W, et al. (2018) Immunological Synapse Predicts Effectiveness of Chimeric Antigen Receptor Cells. Mol Ther 26(4):963–975.

34. Ritter AT, et al. (2015) Actin depletion initiates events leading to granule secretion at the immunological synapse. Immunity 42(5):864–876.

35. Basu R, et al. (2016) Cytotoxic T Cells Use Mechanical Force to Potentiate Target Cell Killing. Cell 165(1):100–110.

36. Tamzalit F, et al. (2019) Interfacial actin protrusions mechanically enhance killing by cytotoxic T cells. Science Immunology 4(33):eaav5445.

37. Kim K, et al. (2016) Three-dimensional label-free imaging and quantification of lipid droplets in live hepatocytes. Scientific Reports 6:36815.

38. Mikolov T, Karafiát M, Burget L, Černocký J, & Khudanpur S (2010) Recurrent neural network based language model. Eleventh annual conference of the international speech communication association.

39. Snoek J, et al. (2015) Scalable bayesian optimization using deep neural networks. International conference on machine learning, pp 2171–2180.

40. Bian C, et al. (2018) Pyramid Network with Online Hard Example Mining for Accurate Left Atrium Segmentation. arXiv e-prints.

41. Chowdhury S, Eldridge WJ, Wax A, & Izatt JA (2017) Structured illumination microscopy for dual-modality 3D sub-diffraction resolution fluorescence and refractive-index reconstruction. Biomed. Opt. Express 8(12):5776–5793.

42. Shin S, Kim D, Kim K, & Park Y (2018) Super-resolution three-dimensional fluorescence and optical diffraction tomography of live cells using structured illumination generated by a digital micromirror device. Scientific Reports 8(1):9183.

43. Kim K, et al. (2017) Correlative three-dimensional fluorescence and refractive index tomography: bridging the gap between molecular specificity and quantitative bioimaging. Biomed. Opt. Express 8(12):5688–5697.

44. Kim K, Yoon J, & Park Y (2015) Simultaneous 3D visualization and position tracking of optically trapped particles using optical diffraction tomography. Optica 2(4):343–346.

45. Kim K & Park Y (2017) Tomographic active optical trapping of arbitrarily shaped objects by exploiting 3D refractive index maps. Nature Communications 8:15340.

46. Rodgers DT, et al. (2016) Switch-mediated activation and retargeting of CAR-T cells for B-cell malignancies. Proceedings of the National Academy of Sciences 113(4):E459.

47. Ronneberger O, Fischer P, & Brox T (2015) U-Net: Convolutional Networks for Biomedical Image Segmentation. Medical Image Computing and Computer-Assisted Intervention – MICCAI 2015, eds Navab N, Hornegger J, Wells WM, & Frangi AF (Springer International Publishing), pp 234–241.

48. Roth HR, et al. (2017) Hierarchical 3D fully convolutional networks for multi-organ segmentation. arXiv e-prints.

49. Dong H, Yang G, Liu F, Mo Y, & Guo Y (2017) Automatic brain tumor detection and segmentation using U-Net based fully convolutional networks. annual conference on medical image understanding and analysis, (Springer), ppp506-517.

50. He K, Zhang X, Ren S, & Sun J (2016) Deep residual learning for image recognition. Proceedings of the IEEE conference on computer vision and pattern recognition, pp 770–778.

51. Peng C, Zhang X, Yu G, Luo G, & Sun J (2017) Large Kernel Matters — Improve Semantic Segmentation by Global Convolutional Network. 2017 IEEE Conference on Computer Vision and Pattern Recognition (CVPR), pp 1743–1751.

