## Supplementary Information for "DeepIS: deep learning framework for three-dimensional label-free tracking of immunological synapses" (from another version of this preprint: version 1, 2019-02-04)

### Deep-learning based three-dimensional label-free tracking and analysis of immunological synapses of chimeric antigen receptor T cells

Moosung Lee[a,b+], Young-Ho Lee[c+], Jinyeop Song [a,b+], Geon Kim[a,b], YoungJu Jo[a,b,d], HyunSeok Min[e], Chan Hyuk Kim[c*], and YongKeun Park[a,b,e*]

[a] Department of Physics, Korea Advanced Institute of Science and Technology (KAIST), Daejeon, South Korea

[b] KAIST Institute for Health Science and Technology, KAIST, Daejeon, South Korea

[c] Department of Biological Sciences, Korea Advanced Institute of Science and Technology, Daejeon, South Korea

[d] Present address: Department of Applied Physics, Stanford University, Stanford, CA 94305, USA

[e] Tomocube Inc., Daejeon, South Korea

+these authors contribute equally to this work.

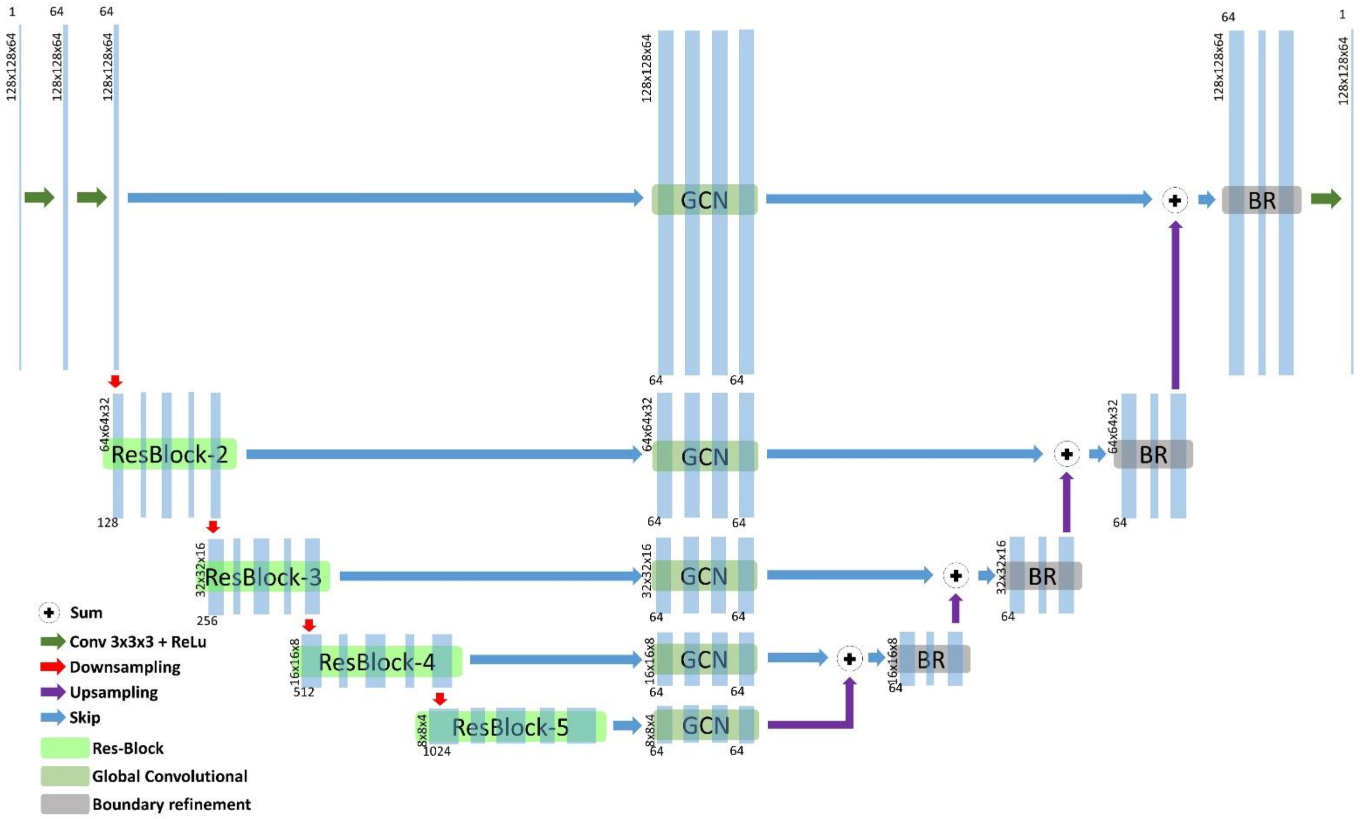


**Supplementary Figure 1. The schematic architecture of our deep neural network.**

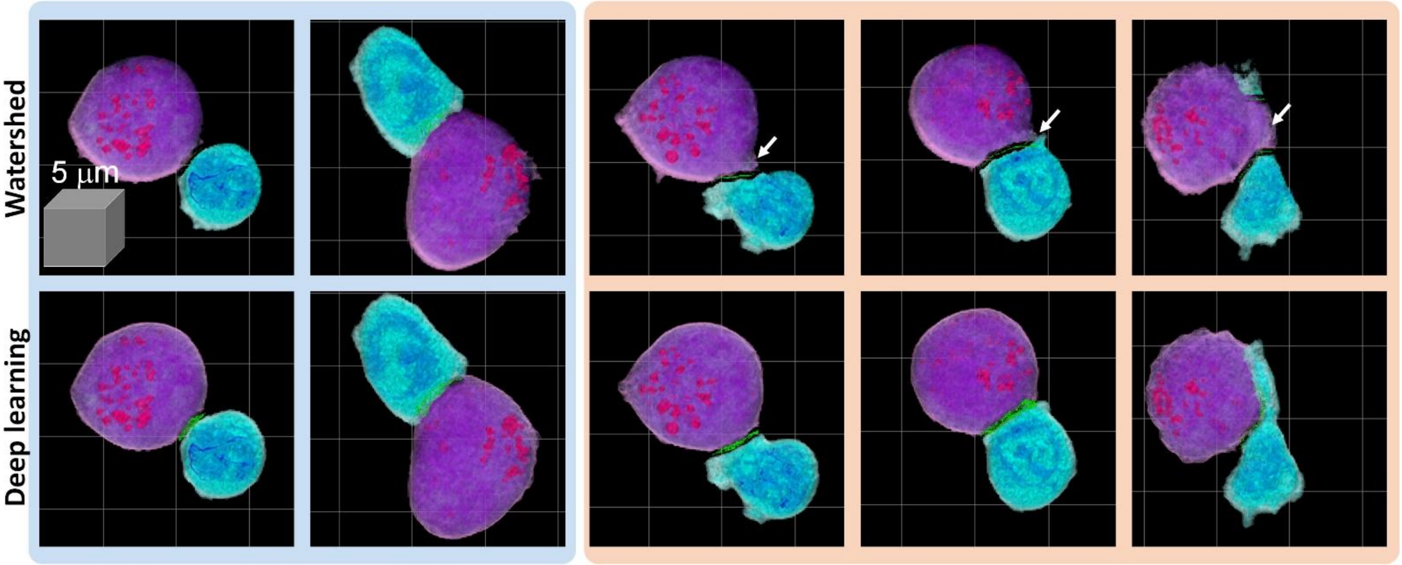


**Supplementary Figure 2. Representative data galleries of segmentation results.** (Top row) Segmentation results using the watershed algorithm. (Bottom row) Segmentation results using deep learning. Blue shade indicates the datasets used in training of deep learning. Red shade indicates the dataset, which was ill-segmented by the watershed algorithm. White arrows illustrate ill-segmented regions.

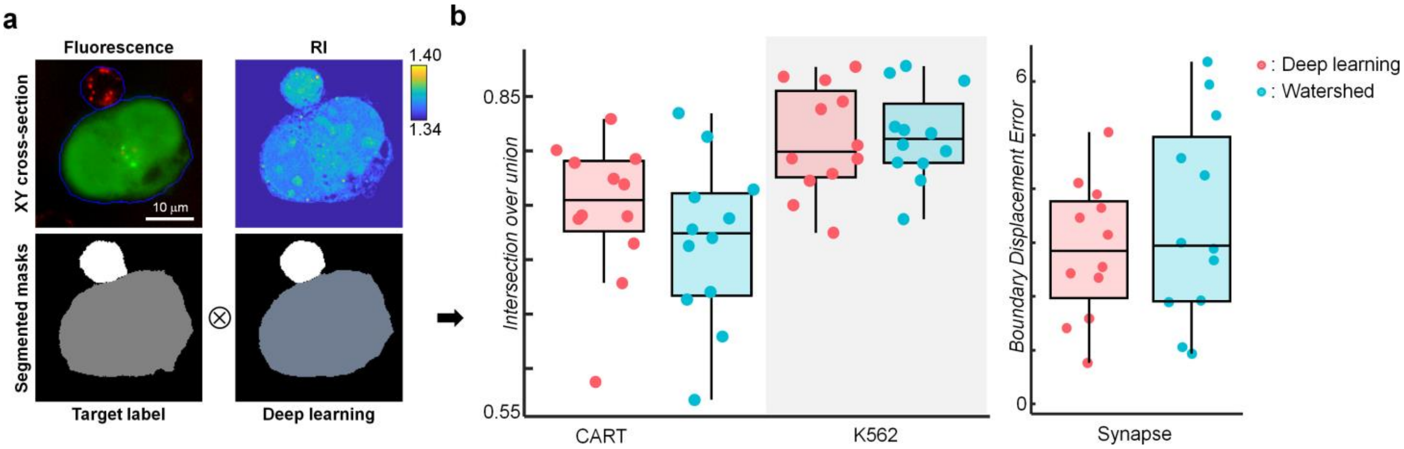


**Supplementary Figure 3. Comparison of segmentation performances between watershed algorithms and our methods. (a)** Representative fluorescence (top left), RI (top right) section images. We obtained labels from RI and FL images semi-manually (bottom left) and compared them with the labels obtained from the trained model (bottom right). **(b)** Intersection over union for CART and K562. (1) CART with Unet (0.746 ± 0.065) (2) CART with watershed (0.717 ± 0.075), (3) K562 with deep learning (0.809 ± 0.049) (4) K562 with deep learning (0.814 ± 0.042). (**c)** Boundary displacement errors for (1) synapse with deep learning (2.86 ± 1.25) (2) synapse with watershed (3.40 ± 1.86). Each boxplot in **b-c** indicates the median, upper, and lower quartiles of each population. The attached lines indicate the range of the population.

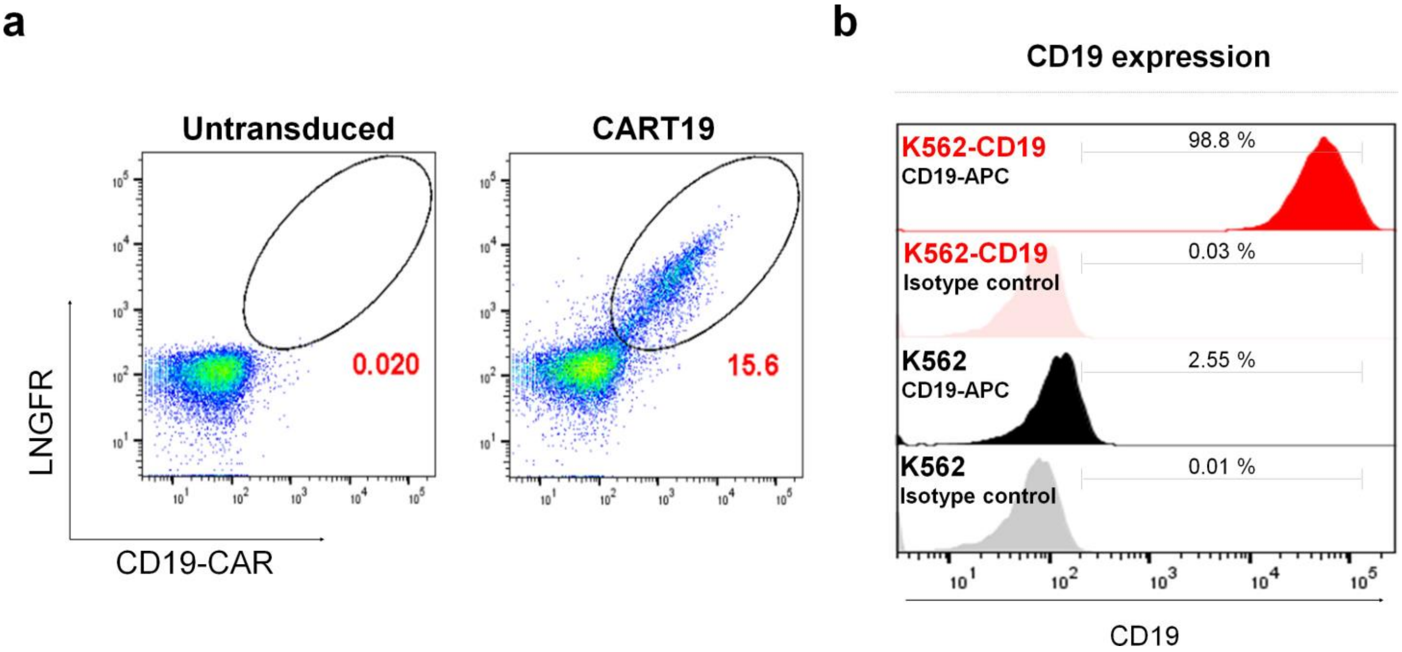


**Supplementary Figure 4. Generations of CD19 specific CAR-T cells and hCD19 transduced target cells for investigating synapse formation dynamics. (a)** LNGFR and CAR expressions at four days after transduction. **(b)** hCD19 expression of K562-CD19 and K562 cells.

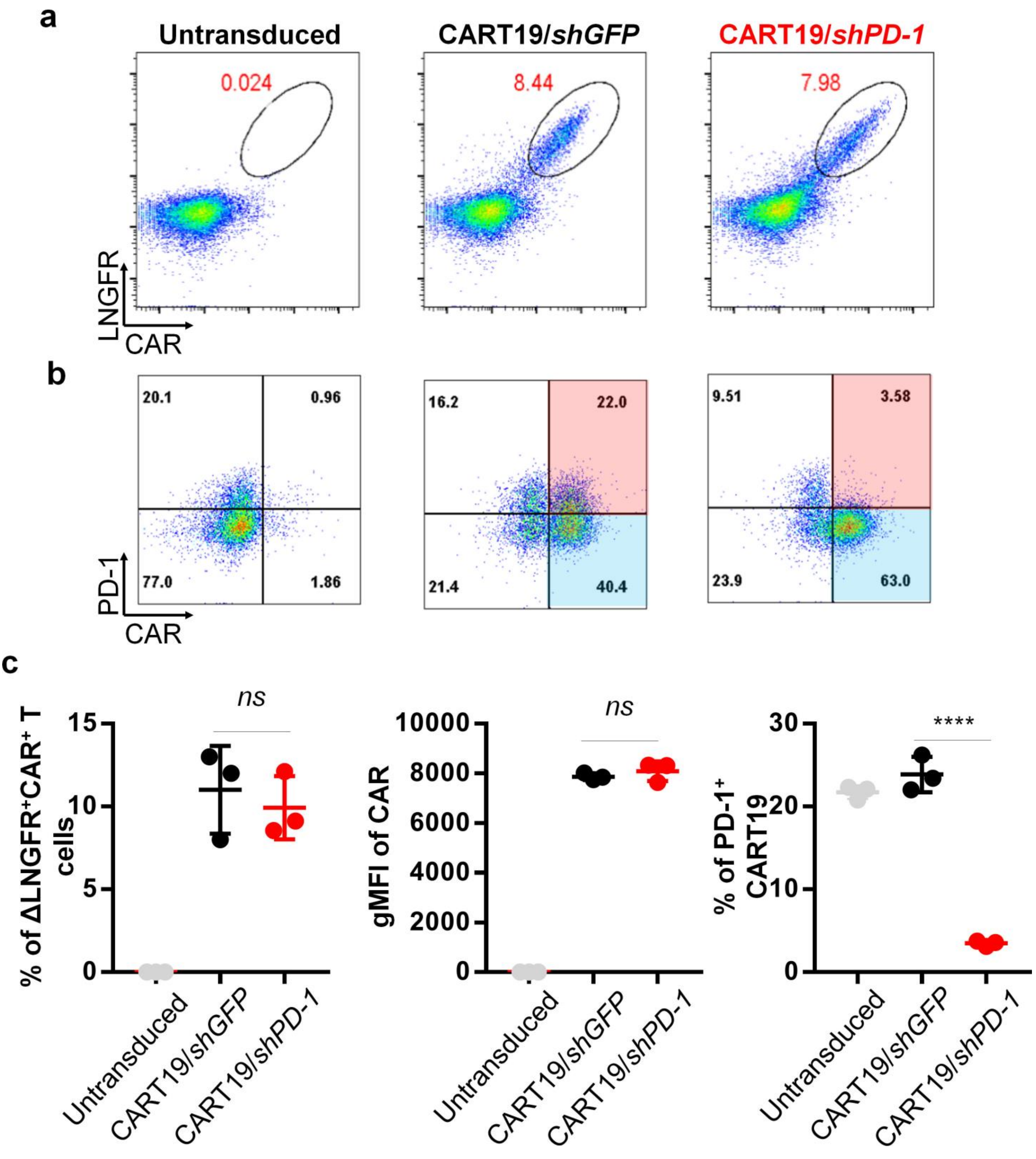


**Supplementary Figure 5. Preparations of untransduced T cells, CART19/*shGFP*, and CART19/*shPD-1* for investigating synapse statistics. (a)** LNGFR and CAR expressions at four days after transduction. Black circles and red figures indicate the transduced cells and their portions in percentages, respectively. **(b)** Surface PD-1 and CD19 specific CAR expressions 72 hours after re-stimulation by mixing CART19 cells with γ-irradiated K562-CD19 without exogenous cytokines. **(c)** Statistics of percentages of LNGFR (left), gMFI of CAR (middle), and percentages of PD-1 positive CART19 (right). ****$p < 0.0001$ by 2-tailed Student`s *t*-test Data represent the mean ± SD.

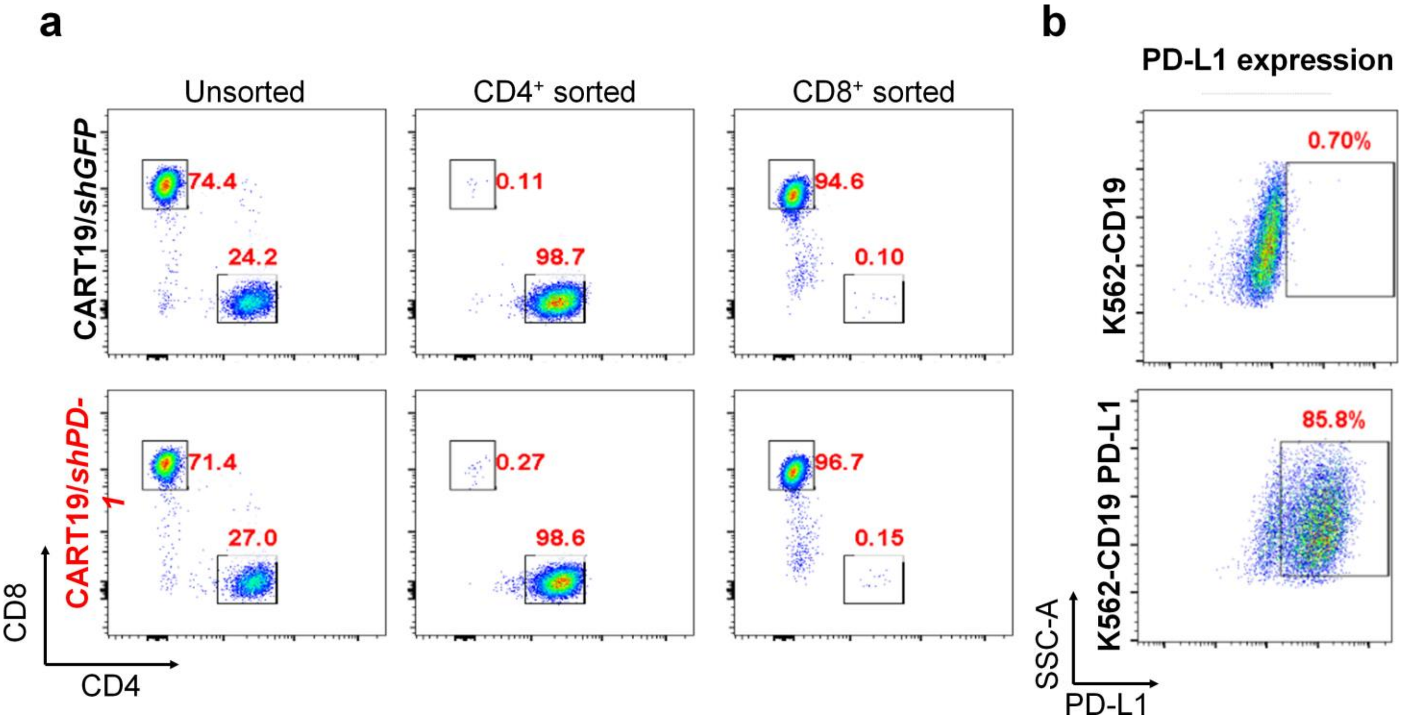


**Supplementary Figure 6. Preparations of CD4$^+$ or CD8$^+$ CART19 cells and K562-CD19-PDL1 cells. (a)** CD4$^+$ or CD8$^+$ CART19 cells were sorted by FACS sorter. **(b)** Establishment of K562-CD19-PDL1 cell line. K562-CD19 cells were transduced by recombinant lentivirus encoding human PD-L1. PD-L1 expression in transduced cells was assessed at four days after transduction. To detection of PD-L1 expression, cells were stained with APC anti-human CD274 (B7-H1, PD-L1) Antibody (Clone: 29E.2A3). PD-L1-transduced K562-CD19 (K562-CD19-PD-L1) cells were sorted for the enrichment of PD-L1$^+$ population.

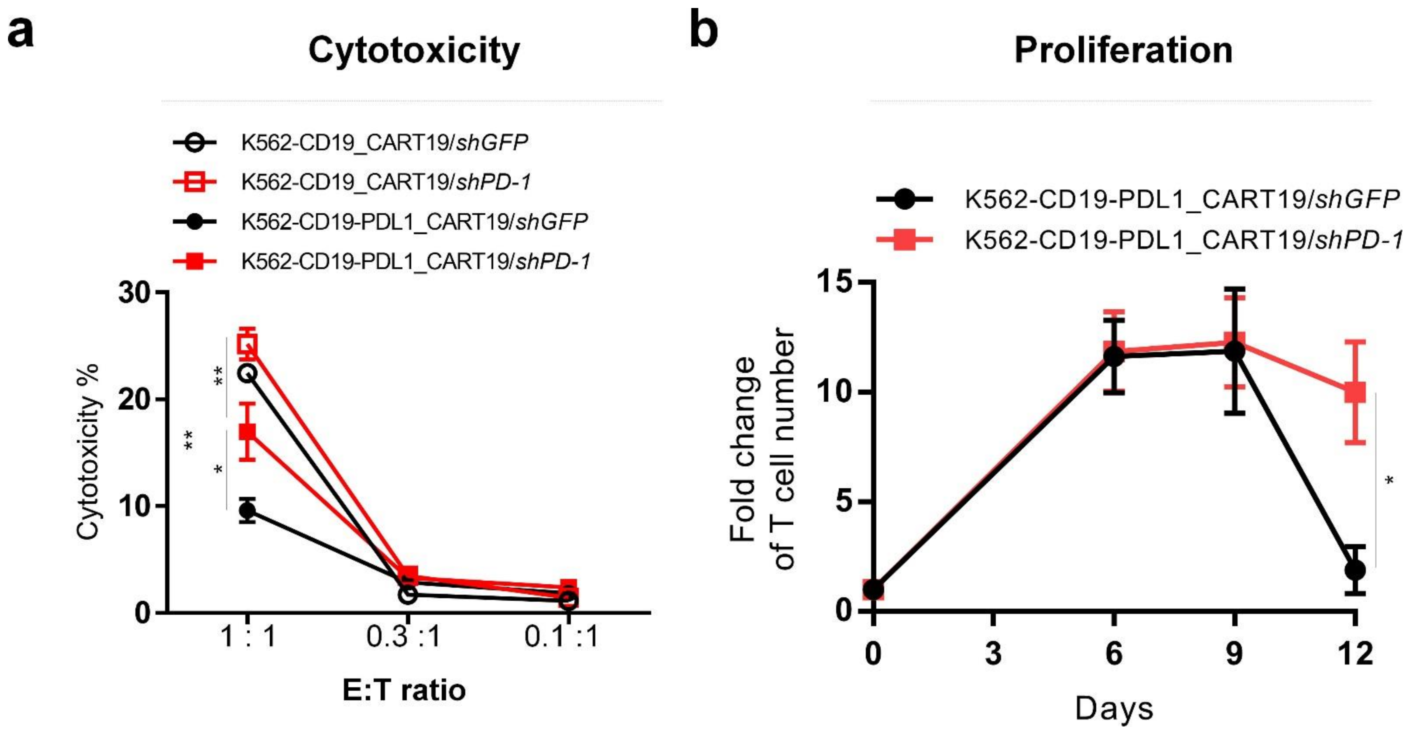


**Supplementary Figure 7. PD-1 knock-down enhanced *in vitro* cytotoxic and proliferative activity under the presence of PD-L1. (a)** LNGFR$^{+}$ CART19 cells were mixed with ClaretRed stained K562-CD19 or K562-CD19-PDL1 at an E: T ratio of 1:1, 0.3:1, and 0.1:1 without exogenous cytokines. Cytotoxicity was assessed 72hr after co-culture with 7-AAD staining. Cytotoxicity % were calculated by 100*(1-(ClaretRed$^{+}$7-AAD$^{-}$ % / ClaretRed$^{+}$ %). **(b)** LNGFR$^{+}$ CART19 cells were mixed with γ-irradiated K562-CD19-PDL1 at an E: T ratio of 1:1. CART19 cell numbers were determined at six days, nine days and 12 days using a trypan blue. Data are representative of at least two independent experiments. *$p < 0.05$, **$p < 0.01$ by 1-way ANOVA (a) or 2-tailed Student`s $t$-test (b). Data represent the mean ± SD.

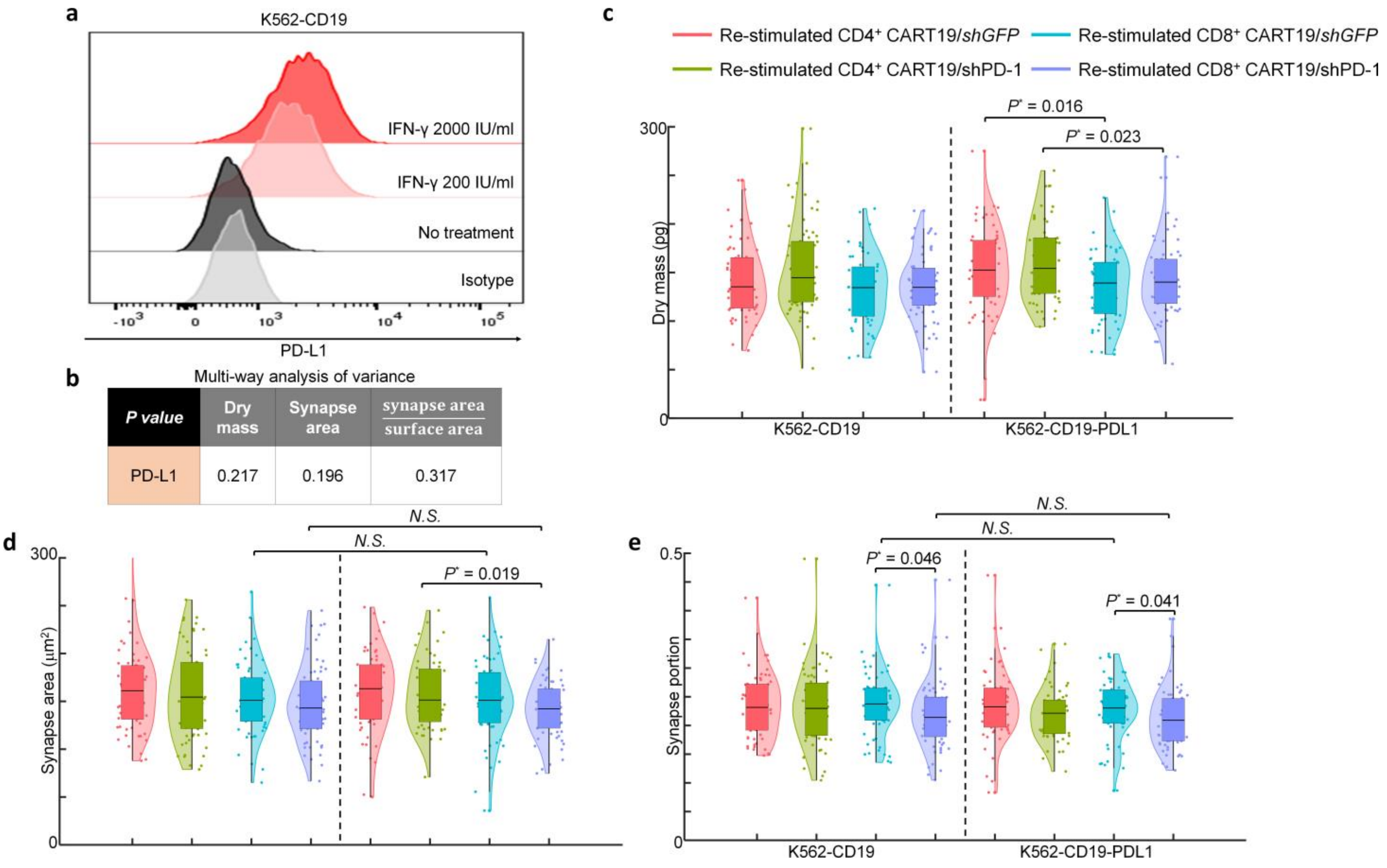


**Supplementary Figure 8. K562-CD19-PDL1 model is not suitable for evaluating the effect of PD-1 / PD-L1 axis on IS. (a)** 24hr PD-L1 expressions of interferon-γ treated K562-CD19 cells. **(b)** Multi-way ANOVA data on PD-L1 expressions of K562-CD19 show the independence of PD-L1 expression levels from either dry mass or synapse area statistics. **(c)** Scatter plots of synapse areas . **(d)** Scatterplots of synapse areas **(e)** Scatterplots of synapse area-per-surface areas $^*$: $p < 0.05$. Mean ± SD are available in Supplementary Table 1.

| Figure | Cell type | Data number | Parameters | mean ± SD | Parameters | mean ± SD | Parameters | mean ± SD |
|---|---|---|---|---|---|---|---|---|
| Figures 3(a-c) | K562 | 707 | Dry mass (pg) | 335.64 ± 109.68 | Synapse area (μm$^2$) | Not available | synapse area-per-surface areas | Not available |
| | resting CD4+ CART19 | 112 | | 58.93 ± 17.70 | | 111.71 ± 29.89 | | 0.294 ± 0.058 |
| | resting CD8+ CART19 | 97 | | 67.35 ± 19.27 | | 113.79 ± 33.88 | | 0.269 ± 0.063 |
| | re-stimulated CD4+ CART19/*shGFP* | 126 | | 147.17 ± 41.15 | | 160.73 ± 43.58 | | 0.235 ± 0.058 |
| | re-stimulated CD4+ CART19/*shPD-1* | 123 | | 156.74 ± 43.68 | | 157.53 ± 41.31 | | 0.226 ± 0.057 |
| | re-stimulated CD8+ CART19/*shGFP* | 122 | | 133.52 ± 36.14 | | 153.45 ± 39.08 | | 0.243 ± 0.124 |
| | re-stimulated CD8+ CART19/*shPD-1* | 127 | | 139.67 ± 36.17 | | 144.81 ± 34.64 | | 0.217 ± 0.057 |
| Supplementary Figures 8(c-e) | re-stimulated CD4+ CART19/*shGFP* with K562-CD19 | 64 | | 142.00 ± 37.48 | | 162.10 ± 43.49 | | 0.236 ± 0.057 |
| | re-stimulated CD4+ CART19/*shPD-1* with K562-CD19 | 67 | | 153.42 ± 46.22 | | 156.83 ± 45.30 | | 0.231 ± 0.065 |
| | re-stimulated CD8+ CART19/*shGFP* with K562-CD19 | 62 | | 132.24 ± 35.91 | | 154.77 ± 36.85 | | 0.257 ± 0.166 |
| | re-stimulated CD8+ CART19/*shPD-1* with K562-CD19 | 65 | | 136.87 ± 33.58 | | 147.96 ± 38.96 | | 0.221 ± 0.061 |
| | re-stimulated CD4+ CART19/*shGFP* with K562-CD19-PDL1 | 62 | | 152.51 ± 44.30 | | 159.33 ± 43.98 | | 0.234 ± 0.061 |
| | re-stimulated CD4+ CART19/*shPD-1* with K562-CD19-PDL1 | 56 | | 160.63 ± 40.51 | | 158.36 ± 36.34 | | 0.219 ± 0.046 |
| | re-stimulated CD8+ CART19/*shGFP* with K562-CD19-PDL1 | 60 | | 134.85 ± 36.63 | | 152.09 ± 41.52 | | 0.228 ± 0.053 |
| | re-stimulated CD8+ CART19/*shPD-1* with K562-CD19-PDL1 | 62 | | 142.60 ± 38.77 | | 141.52 ± 29.40 | | 0.213 ± 0.053 |

**Supplementary Table 1. Mean and standard deviations of IS-related parameters.**
